# Rad27/FEN1 prevents accumulation of unprocessed Okazaki fragments and ribosomal DNA copy number changes

**DOI:** 10.1101/2025.03.10.642467

**Authors:** Tsugumi Yamaji, Yuko Katayama, Nanase Arata, Mariko Sasaki

## Abstract

DNA copy number changes are the most frequent genomic alterations in cancer cells. The ribosomal DNA (rDNA) region is particularly vulnerable to such changes due to its repetitive nature. Here, we demonstrate that Rad27/FEN-1, a structure-specific nuclease in budding yeast, plays a crucial role in maintaining rDNA stability. The production of extrachromosomal rDNA circles and severe chromosomal rDNA instability are observed in the *rad27*Δ mutant, independently of Fob1-mediated DNA replication fork arrest and DNA double-strand break (DSB) formation in the rDNA. The *rad27*Δ mutant accumulates unprocessed Okazaki fragments in the rDNA region, without inducing DSB formation. Similar rDNA instability is observed in DNA ligase^Cdc9^-deficient cells. Furthermore, we show that Exonuclease 1 and PCNA can compensate for the loss of Rad27 function in the rDNA stabilization. These findings highlight the importance of proper Okazaki fragment processing in preventing non-DSB-induced rDNA copy number changes.

## INTRODUCTION

DNA copy number changes are among the most frequent genomic alterations observed in cancer cells (*1, 2*). These changes can occur through deletion or amplification of focal DNA segments on chromosomes, accumulation of extrachromosomal circular DNAs, or gain or loss of entire chromosomes (*3-6*). Such changes often lead to altered gene expression, driving tumorigenesis and other diseases (*5-7*).

The ribosomal RNA gene (rDNA) is essential for ribosome biogenesis and is present in high copy number in eukaryotic genomes, forming one or several clusters of tandemly arrayed rDNA copies. In budding yeast, approximately 150 rDNA copies are arrayed at a single locus on chr XII (Fig. 1A). The repetitive nature of rDNA clusters makes them prone to amplification or deletion, as well as the formation of extrachromosomal rDNA circles (ERCs). To counter this instability, organisms have evolved mechanisms to maintain rDNA copy number (*8-10*), which is best understood in budding yeast. After DNA replication is initiated from an origin of DNA replication, two replication forks proceed bi-directionally but the fork moving in the opposite direction to the 35S rDNA is blocked by Fob1 bound to the replication fork barrier (RFB) site, resulting in a single-ended DSB (Fig. 1A) (*11-16*). Repair of these DSBs can result in changes in the chromosomal rDNA copy number and is often associated with ERC production (Fig. 1A) (*10*). Many mutants defective in rDNA stability have been identified, and for genes whose functions have been characterized in details, many primarily act in the Fob1-dependent pathway (*17-21*).

**Figure 1.**
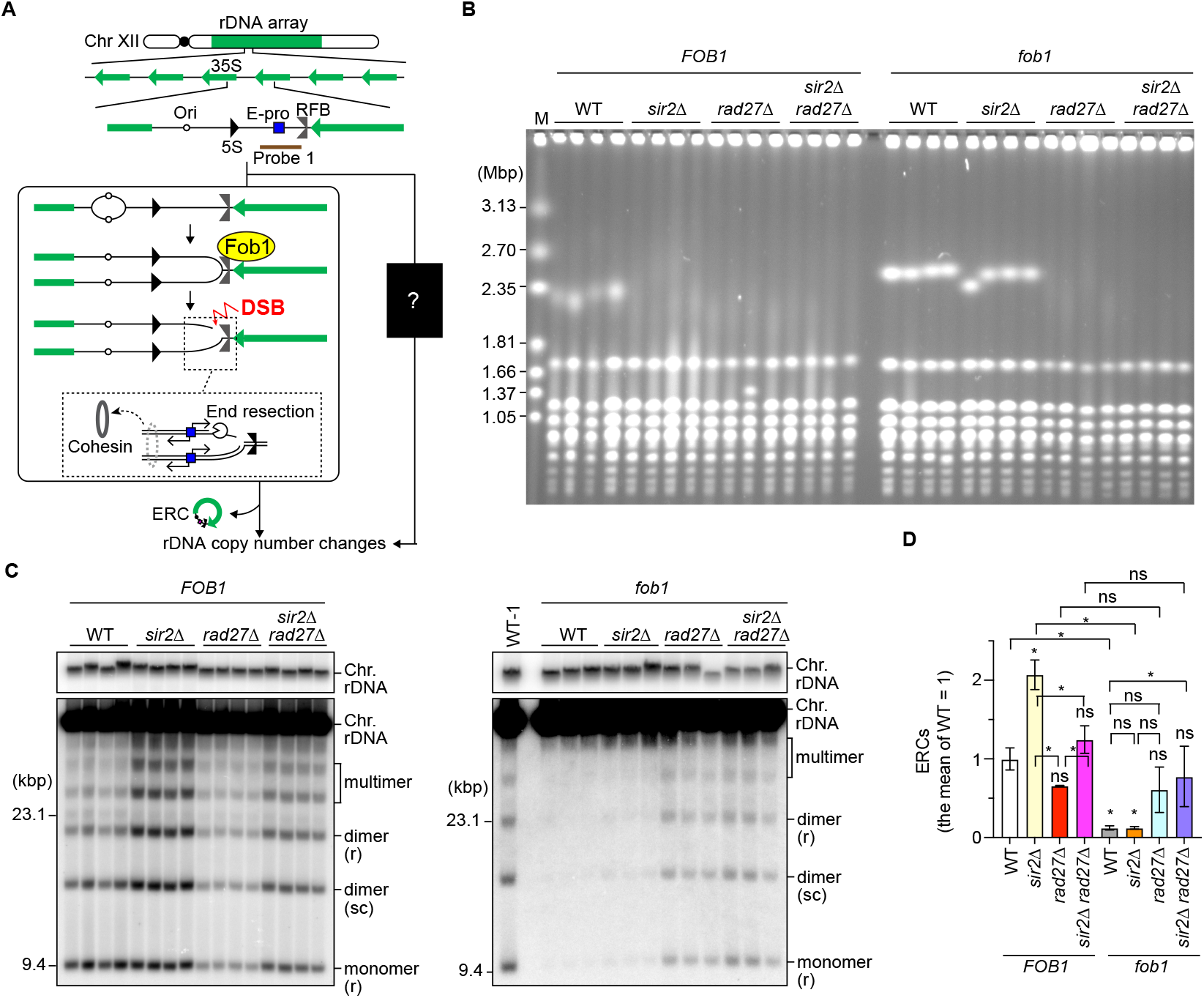
Rad27 stabilizes the rDNA in a Fob1-independent manner. **(A)** The rDNA cluster in the budding yeast genome. Budding yeast carries a single rDNA cluster on Chr XII, where approximately 150 rDNA copies are tandemly arrayed. Each copy contains 35S rRNA (35S), 5S rRNA (5S), an origin of DNA replication (Ori), a bidirectional promoter for noncoding RNA, called E-pro, and a replication fork barrier (RFB). The left panel shows the previously known Fob1-dependent pathway that leads to chromosomal rDNA copy number changes and the production of ERCs. Probe 1 was used for the ERC assay. **(B)** PFGE analysis to examine the size heterogeneity of chr XII. DNA was isolated from four independent clones of the indicated strains, separated by PFGE, and stained with ethidium bromide (EtBr). M indicates *Hansenula wingei* chromosomal DNA markers. **(C)** ERC detection. DNA was isolated from four independent clones of WT, *sir2*Δ, *rad27*Δ, and *sir2*Δ *rad27*Δ and three independent clones of *fob1, fob1 sir2*Δ, *fob1 rad27*Δ, and *fob1 sir2*Δ *rad27*Δ. DNA was separated by agarose gel electrophoresis, followed by Southern blotting with probe 1, as shown in (A). On the right gel, DNA from the first WT clone run on the left gel was also loaded as WT-1. Chromosomal rDNA and different forms of ERCs are indicated. r and sc indicate relaxed and supercoiled forms of ERCs, respectively. Supercoiled monomers ran off from the gel under the electrophoresis conditions used. M indicates λ DNA-Hind III markers. Top panels show short exposure of genomic rDNA signals. **(D)** The level of ERCs. The sum of monomers, dimers, and multimers was normalized to the chromosomal rDNA signals. The level of ERCs was normalized to the average of the WT clones. Bars show the mean ± the standard error of the mean (s.e.m.). The difference between strains was compared by one-way ANOVA, followed by Tukey’s multiple comparisons test. Asterisks above the bars and brackets indicate statistically significant differences (P < 0.05) between the WT strain and the indicated strains and between different strains, respectively. ns indicates no statistically siginificant difference.

During DNA replication, lagging strand synthesis occurs discontinuously (*22, 23*). DNA polymerase α-primase initiates the process by generating RNA-DNA primers. These primers are then extended by DNA polymerase δ, with the assistance of Proliferating Cell Nuclear Antigen (PCNA). Upon encountering a previously synthesized Okazaki fragment, DNA polymerase δ displaces the RNA-DNA primer, generating 5’ flap structures. These flaps are cleaved by nucleases including the 5’ flap endo/exonuclease Rad27 (FEN1 in mammals), Dna2, Exo1, and RNase H (*22, 24*). FEN1 deficiency causes genomic instability such as accumulation of somatic mutations and short repeat instability and cancer predisposition (*25*). Finally, Okazaki fragments are joined by DNA ligase Cdc9.

In this study, we demonstrate that Rad27 is crucial for maintaining rDNA stability. Unlike previously characterized rDNA-stabilizing factors, Rad27 functions independently of the Fob1-dependent pathway, which responds to programmed replication fork arrest and DSBs at the RFB. By conducting physical assays, we demonstrate that absence of Rad27 leads to the accumulation of unprocessed Okazaki fragments in the rDNA *in vivo*, without detectable DSB formation. The temperature-sensitive *cdc9-1* mutant also shows severe rDNA instability, similar to the *rad27*Δ mutant. Finally, overexpression of *EXO1* and PCNA^*POL30*^ alleviates rDNA instability in *rad27*Δ cells. We propose that Rad27-mediated processing of Okazaki fragments is crucial for rDNA stabilization.

## RESULTS

### Rad27 stabilizes the rDNA by preventing the occurrence of Fob1-independent lesions

A previous genome-wide screen identified that cells lacking Rad27 exhibit chromosomal rDNA instability (*21*). We confirmed this phenotype and compared the degree of rDNA instability in the *rad27*Δ mutant to that of the *sir2*Δ mutant, a known rDNA instability mutant (*15, 26*). Genomic DNA was isolated from these mutants, separated it by pulsed-field gel electrophoresis (PFGE), and the size heterogeneity of chr XII carrying the rDNA array was examined. The wild-type (WT) strain displayed homogeneous chr XII bands, while the *sir2*Δ mutant showed extremely smeared bands, indicating frequent changes in chromosomal rDNA copy number in the *sir2*Δ mutant (Fig. 1B). The *rad27*Δ mutant also showed a similar degree of smearing of chr XII bands as the *sir2*Δ mutant (Fig. 1B). Thus, Rad27 deficiency results in severe rDNA instability by causing both contraction and expansion of the chromosomal rDNA array.

Previously characterized rDNA-unstable mutants show increased ERC levels along with chromosomal rDNA copy number changes (*17, 20, 26, 27*). Consistent with these findings (*20, 21, 26*), the ERC level in the *sir2*Δ mutant was >2-fold higher than that in WT cells (Fig. 1C). On the contrary, the level of ERCs in the *rad27*Δ mutant was similar to WT cells, demonstrating that chromosomal rDNA instability in the *rad27*Δ mutant was not accompanied by enhanced ERC production in the WT (*FOB1*) background.

Because Sir2 promotes accurate repair of DSBs at Fob1-dependent arrested forks, the *fob1 sir2*Δ mutant showed sharp chr XII bands and low ERC levels, similar to those seen in the *fob1* single mutant. In contrast, an introduction of a *fob1* mutation did neither affect the degree of smearing of chr XII bands nor ERC levels in the *rad27*Δ mutant (Fig. 1C–1F). These findings suggest that Rad27 prevents Fob1-independent lesions that trigger rDNA instability.

### Interaction of Rad27 with PCNA is important for the maintenance of rDNA stability

Rad27 is a member of the Rad2/XPG family of structure-specific nucleases (*28*). We examined whether other nucleases in this family are also required for the maintenance of rDNA stability. Unlike the *rad27*Δ mutant, *din7*Δ, *yen1*Δ, *exo1*Δ, and *rad2*Δ mutants showed sharp chr XII bands, comparable to those in the WT strain (Fig. 2A). It is possible that these nucleases function redundantly with Rad27. However, Rad27 plays a major role in rDNA stabilization.

**Figure 2.**
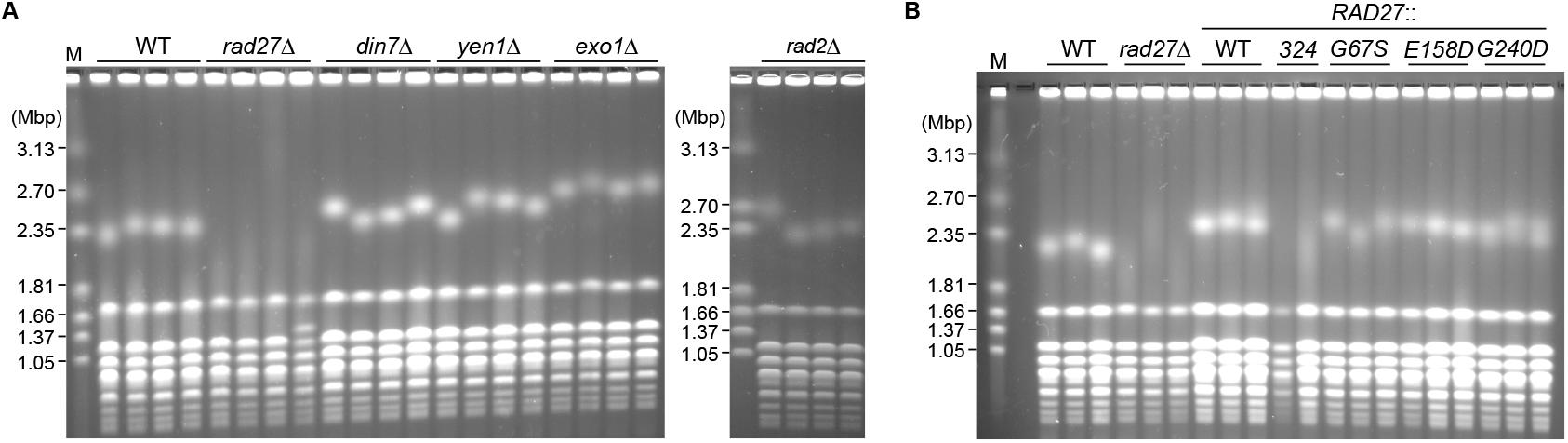
Interaction of Rad27 with PCNA is important for the maintenance of rDNA stability. **(A, B)** PFGE analysis to examine the size heterogeneity of chr XII. In (A), DNA isolated from four independent clones of the indicated strains was analyzed. In (B), DNA isolated from three independent clones of WT and *rad27*Δ strains was analyzed on the leftmost lanes, and DNA isolated from strains where the endogenous *RAD27* allel was replaced with WT or indicated *rad27* alleles was analyzed. DNA was separated by PFGE and stained with EtBr. M indicates *H. wingei* chromosomal DNA markers.

To investigate how Rad27 stabilizes the rDNA, we used a series of *rad27* mutant alleles defective in different activities or properties. The *rad27-324\** mutation introduces a premature stop codon at amino acid 324, resulting in a truncated protein lacking the C-terminal PCNA-interacting protein box (*29*). When the endogenous *RAD27* was replaced with the *rad27-324\** allele, cells showed smeared chr XII bands, similar to those seen in the *rad27*Δ mutant (Fig. 2B), suggesting that Rad27’s interaction with PCNA is important for preventing rDNA copy number changes.

The *rad27-G67S* and *rad27-G240D* alleles carry mutations in the N and I nuclease domains, respectively. *In vitro*, the rad27-G67S protein showed reduced activity in cleaving single 5’-end flap substrates, while the rad27-G240D protein is severed impaired in this activity (*29*). However, both rad27-G67S and rad27-G240D proteins can cleave double-flap substrates *in vitro* (*29*), which contain a 3’ flap adjacent to the 5’ flap and are thought to be preferred substrates for Rad27 *in vivo* (*24*). The *rad27-G67S* and *rad27-G240D* mutants showed homogeneous chr XII bands, comparable to those in WT cells (Fig. 2B). Thus, the activity to cleave a single 5’ flap is not essential for rDNA stabilization. Lastly, the *rad27-E158D* mutation, located in the active site of Rad27 at aspartate 158, severely compromises its exonuclease activity, while retaining partial endonuclease activity toward the 5’ flap substrate (*30*). The *rad27-E158D* mutant showed homogeneous chr XII bands, indicating that the Rad27’s exonuclease activity is dispensable for rDNA stabilization (Fig. 2B). Currently, mutations that specifically eliminate the activity to cleave double flap substrates have not been identified, so its importance for rDNA stabilization cannot be assessed. These findings suggest that the interaction of Rad27 with PCNA is crucial for maintaining rDNA stability.

### Rad27 facilitates transcription from E-pro in the *sir2*Δ mutant background

Transcription from E-pro can trigger rDNA instability in the Fob1-dependent pathway by inducing end resection of replication-coupled DSBs formed at the RFB and also by inhibiting cohesin association with the rDNA (*15, 26, 31*). Rad27 has been shown to suppress the accumulation of telomere repeat-containing RNA (TERRA) (*32*). Therefore, we examined whether Rad27 is involved in regulating transcription from E-pro. To this end, total RNA was isolated, separated by agarose gel electrophoresis under denaturing conditions, and non-coding RNAs from E-pro was detected by Northern blotting with strand-specific probes (Fig. 3A).

**Figure 3.**
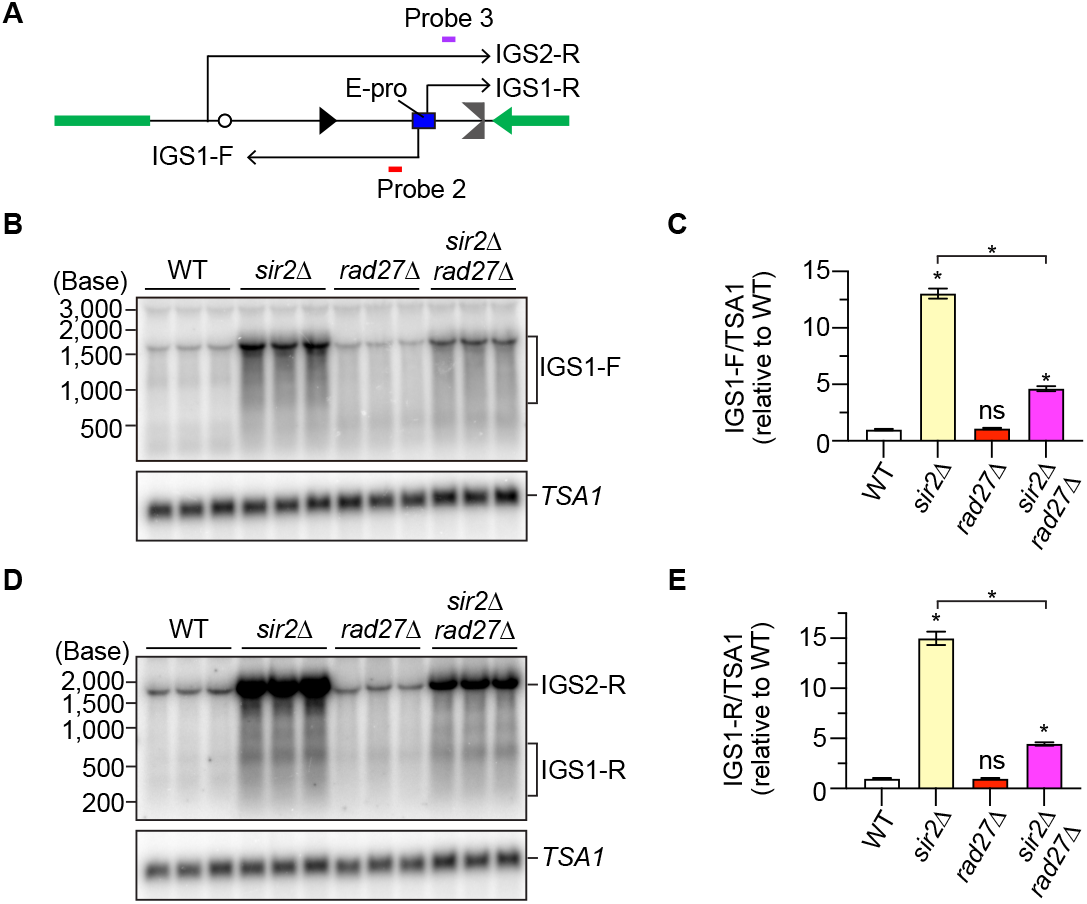
Rad27 facilitates transcription from E-pro in the *sir2*Δ mutant. **(A)** Transcription of non-coding RNA in the rDNA. Probes 2 and 3 are single-stranded DNA probes used to detect IGS1-F and IGF1-R transcribed from the E-pro, respectively. **(B, D)** Detection of non-coding RNAs. IGS1-F (B) and IGS1-R (D) were detected by Northern blotting with the probes indicated in (A). The same membrane was re-probed to detect *TSA1* transcripts. **(C, E)** Levels of non-coding RNA transcripts. The level of IGS1-F (C) and IGS1-R (E) was quantified relative to that of *TSA1*, which was normalized to the average level of transcripts in the WT. Bars show the mean ± s.e.m. The difference between strains was compared by one-way ANOVA, followed by Tukey’s multiple comparisons test. Asteriks above the bars and brackets indicate statistically significant differences (P < 0.05) between the WT strain and the indicated strains and between different strains, respectively. ns indicates no statistically significant difference.

IGS1-R transcripts of ∼300–650 nt in size were observed in the *sir2*Δ mutant, where transcription from E-pro is derepressed (Fig. 3B), consistent with previous findings (*17, 33, 34*). The level of these transcripts was ∼15-fold higher in the *sir2*Δ mutant than that in WT (Fig. 3C). Deletion of *RAD27* did not affect IGS1-R transcript levels (Fig. 3C). However, while deletion of *RAD27* in the WT strain did not affect their transcript level, its deletion in the *sir2*Δ mutant caused a reduction in the level of IGS1-R transcripts (Fig. 3B, C). IGS1-F transcripts of the size ranging from ∼800 to ∼1,500 nt were detected in the *sir2*Δ mutant at the level >10-fold higher than that in WT cells (Fig. 3D, E), consistent with previous findings (*17, 33, 34*). The *rad27*Δ mutant showed IGS1-F transcripts at the level comparable to that in WT cells. Similar to IGS1-R transcripts, deletion of *RAD27* in the *sir2*Δ mutant resulted in a 3-fold decrease in IGS1-F transcript levels (Fig. 3D, E). Taken together, these findings indicate that Rad27 has less impact on transcription from E-pro in the WT strain but promotes transcription of noncoding RNA in the *sir2*Δ mutant.

### Absence of Rad27 accumulates unprocessed Okazaki fragments in the rDNA

Rad27 is involved in Okazaki fragment maturation and has also been implicated in other pathways such as mismatch repair (*35*). To determine which function is important for rDNA stabilization, we examined the phenotype of a mutant lacking Cdc9, the ligase that joins Okazaki fragments (*36*). We isolated temperature-sensitive *cdc9-1* clones, grew them at the semi-permissive temperature, and analyzed their rDNA instability. The *cdc9-1* mutant showed extremely smeared chr XII bands, similar to those seen in the *rad27*Δ mutant (Fig. 4A).

**Figure 4.**
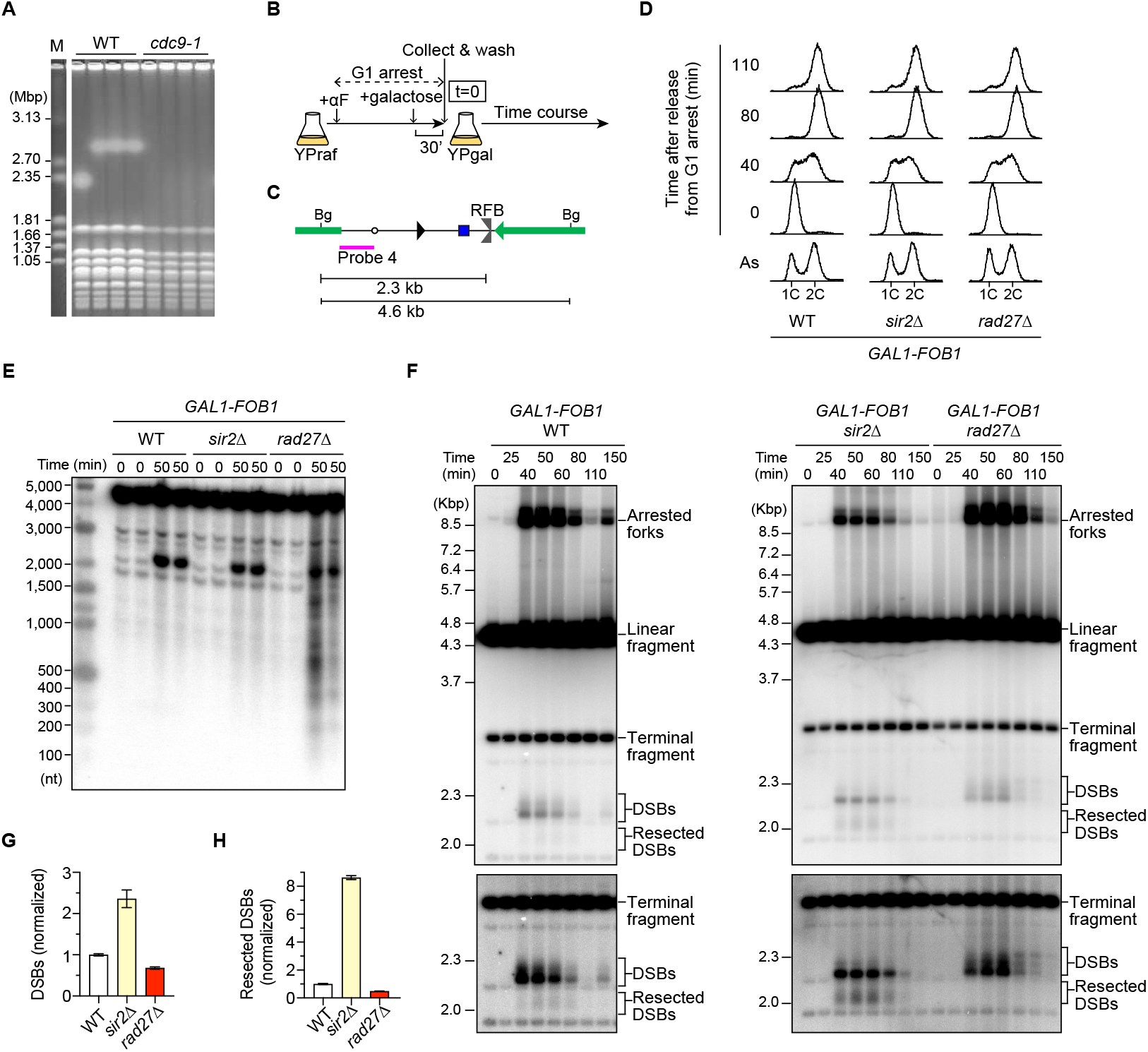
The *rad27*Δ mutant accumulates Okazaki fragments in the rDNA. **(A)** PFGE analysis to examine the size heterogeneity of chr XII. Four independent clones of WT and the temperature sensitive *cdc9-1* mutant were grwon at the semi-permissive temperature. DNA was isolated, separated by PFGE, and stained with EtBr. M indicates *H. wingei* chromosomal DNA markers. **(B)** The outline of the time course experiment. **(C)** The position of probe 4 used for SSB and DSB assays. *Bg* indicates a recognition site for the restriction enzyme Bgl II. **(D)** Cell cycle analysis of time courses in the indicated strains by flow cytometry. **(E)** SSB assay. DNA was isolated from the indicated strains collected at t=0 and 50 min after release from G1 arrest in two independent time course experiments. DNA was separated by denaturing agarose gel electrophoresis, followed by Southern blotting with probe 4. **(F)** DSB assay. Arrested forks, linear fragments, DSBs, and resected DSBs are indicated. The terminal fragment corresponds to the restriction fragment at the telomere-proximal rDNA repeat. Bottom panels show long exposure images, only showing the region around DSBs and resected DSBs. **(G, H)** Quantitation of DSBs and resected DSBs at the RFB. The frequency of DSBs (G) and resected DSBs (H) was estimated by calculating the ratio of the DSB signal relative to that of arrested forks in (G) and the ratio of the resected DSB signal relative to that of DSBs in (H), which was normalized to the average of WT clones. Bars show the range from two independent experiments.

To examine whether the *rad27*Δ mutant accumulates single-strand DNA breaks (SSBs) in the rDNA region, we arrested cells in G1 phase and collected cells immediately after release from G1 arrest (Fig. 4B). We also collected cells 50 min after release from G1 arrest, when most cells were in S phase. Genomic DNA was prepared, digested with Bgl II, and separated DNA by agarose gel electrophoresis under denaturing conditions, followed by Southern blotting with an rDNA probe (Fig. 4C). Prominent signals observed in WT and the *sir2*Δ mutant showed S-phase specific signals at >2 kb, which corresponds to strands composed of DSBs at the RFB (Fig. 4D). In addition to the DSB signals, the *rad27*Δ mutant showed S-phase specific signals throughout the region analyzed. Okazaki fragment synthesis is interlinked with chromatin assembly, and the length of lagging strands is influenced by nucleosomes (*37*). The *rad27*Δ mutant showed ladders of signals that appeared to match nucleosome sizes (Fig. 4D). Therefore, Rad27 prevents the accumulation of unprocessed Okazaki fragments in the rDNA.

Deletion of *RAD27* confers synthetic lethality with mutations in *RAD52*, an essential protein for homologous recombination (HR)-mediated DSB repair (*38, 39*). Thus, unprocessed Okazaki fragments may be converted into DSBs, inducing Rad52-mediated repair and subsequent rDNA instability. To test this possibility, we performed time course experiments during synchronous S phase progression, prepared genomic DNA, digested it with Bgl II (Fig. 4B), and separated it by conventional agarose gel electrophoresis, followed by Southern blotting (Fig. 4C). DSBs were formed at the RFB during S phase in all strains examined. In the *sir2*Δ mutant, DSBs were formed more frequently at the RFB, and these DSB ends were resected more frequently, compared to WT cells, inducing HR-mediated rDNA instability, consistent with a previous finding (*26*). In contrast, the absence of Rad27 affected neither the level of DSBs at the RFB nor the frequency of its end resection, indicating that Rad27 is unlikely to impact DSB formation and its repair at the RFB. Importantly, Rad27 deficiency did not lead to the formation of DSBs at sites other than the RFB. Taken together, these findings suggest that unprocessed Okazaki fragments in the *rad27*Δ mutant are the likely trigger for rDNA instability.

### Exo1 and Pol30 compensate for the function of Rad27 in rDNA stabilization

Several factors have been implicated in Okazaki fragment maturation (*22, 23, 38*). To investigate whether overexpression of any of these factors can compensate for Rad27’s function, we constructed plasmids carrying *DNA2, EXO1, PIF1, RNH201, POL30*, and *CDC9*, introduced them into *rad27*Δ cells, and analyzed their impact on rDNA stability. The YCp22 plasmid is an ARS/CEN plasmid, which is maintained in yeast cells at a copy of ∼1–2 copies per cell. Introduction of an empty vector in the *rad27*Δ mutant showed smeared chr XII bands, but expression of *RAD27* restored sharp chr XII bands (Fig. 5A). Overexpression of *EXO1* in the *rad27*Δ mutant sharpened the chr XII bands, indicating that Exo1 can compensate for Rad27 in stabilizing the rDNA. On the contrary, overexpression of *DNA2, PIF1, RNH201*, and *CDC9* did not cause sharpening of chr XII bands. Overexpression of *POL30* led to slightly sharper chr XII bands, compared to the *rad27*Δ mutant with an empty vector, indicating that *POL30* can partially suppress rDNA instability caused by *rad27*Δ (Fig. 6A). These findings indicate that Exo1 and PCNA^Pol30^ can facilitate rDNA stabilization in the absence of Rad27.

**Figure 5.**
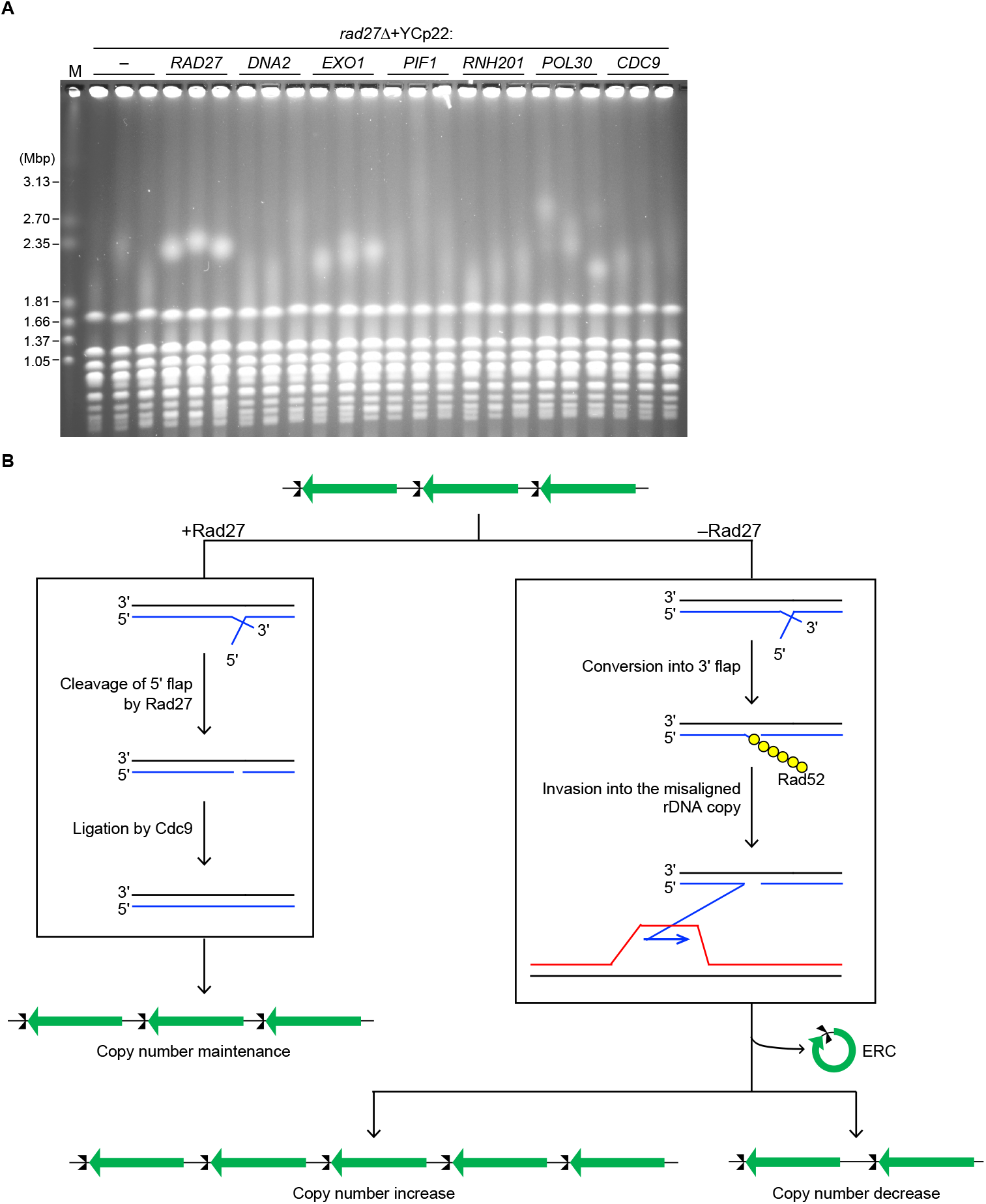
Overexpression of *EXO1* and *POL30* suppresses rDNA instability of the *rad27*Δ mutant. **(A)** PFGE analysis to examine the size heterogeneity of chr XII. DNA was isolated from three independent clones of the *rad27*Δ mutant carrying an empty vector or the plasmid carrying the indicated genes. DNA was separated by PFGE and stained with EtBr. M indicates *H. wingei* chromosomal DNA markers. **(B)** Model for rDNA copy number changes induced by unprocessed Okazaki fragments. In the presence of Rad27, Okazaki fragments are properly matured by clavage of 5’ flap structures, followed by ligation by a Cdc9 DNA ligase, leading to the rDNA copy number maintenance. In the absence of Rad27, Okazaki fragment maturation is inefficient due to the accumulation of 5’ flap structures. The 5’ flap can be converted into a 3’ flap, to which Rad52 binds and carries out strand invasion into a misaligned rDNA copy, leading to non-DSB-induced rDNA copy number changes and ERC production.

## DISCUSSIONS

In this study, we demonstrate that the mutant lacking Rad27/FEN-1 and the temperature-sensitive DNA ligase *cdc9-1* mutant display severe rDNA instability in a Fob1-independent manner. We show that the *rad27*Δ mutant accumulates S-phase-specific SSBs, derived from unprocessed Okazaki fragments, although distinct DSBs are not evident specifically in *rad27*Δ cells. Our genetic data further demonstrate that the other nuclease Exo1, but not Dna2, can substitute for the role of Rad27 in rDNA stabilization.

Previously characterized rDNA-unstable mutants exhibit Fob1-dependent rDNA instability, although the degree of dependency varies (*17, 21*). For example, rDNA instability in the *sir2*Δ mutant is primarily Fob1-dependent (*15, 26*), while rDNA instability in the mutant lacking the histone chaperone Asf1 is partially Fob1-dependent (*18*). On the contrary, the *rad27*Δ mutant displayed severe chromosomal rDNA instability that was not suppressed by an introduction of a *fob1* mutation (Fig. 1B, 1C). Furthermore, while ERC production is suppressed by *fob1* in other rDNA-unstable mutants (*17, 21, 26*), ERC levels were comparable in *rad27*Δ and *fob1 rad27*Δ mutants (Fig. 1D, 1E). There findings suggest that in cells lacking Rad27, lesions other than Fob1-dependent DSBs at the arrested replication forks trigger rDNA instability.

Although Rad27 deficiency did not affect E-pro transcript levels in the WT background, it led to a reduction in their levels in the *sir2*Δ mutant background (Fig. 3). This suggests that Rad27 can promote transcription from E-pro, which contrasts with its role in suppressing the accumulation of TERRA in the telomeric region (*32*). During S phase, RNA polymerase can use both the lagging strand and the template for the leading strand as a template for RNA synthesis. Because Okazaki fragments are prone to remain unprocessed and unligated in the absence of Rad27 (Fig. 4E), we speculate that the level of noncoding RNA from E-pro was reduced due to inefficient transcription on the lagging strand. This effect is more evident in the *sir2*Δ mutant, where transcription from E-pro is derepressed (Fig. 3).

We demonstrate that S-phase cells lacking Rad27 accumulate SSBs, reminiscent of unprocessed Okazaki fragments (Fig. 4E). Furthermore, the *cdc9-1* mutant displays chromosomal rDNA instability comparable to that seen in *rad27*Δ cells (Fig. 4A). These findings suggest that unprocessed Okazaki fragments are responsible for rDNA instability.

Rad27, Dna2, and Exo1 have been implicated in processing 5’ flaps generated during Okazaki fragment synthesis, based on genetic and biochemical analyses (*38, 40-42*). Kahli et al, demonstrated that *in vivo*, Rad27 is the primary enzyme that cleaves lagging-strand flaps, and that Exo1, but not Dna2, can compensate for its absence, by conditionally depleting these nucleases alone or in combinations and then analyzing lagging strands by physical assays (*43*). Our finding that overexpression of *EXO1*, but not of *DNA2*, suppressed rDNA instability in the *rad27*Δ mutant (Fig. 5A) supports these findings (*43*). Interestingly, the absence of Exo1 alone did not result in rDNA instability (Fig. 2A), suggesting that Rad27 plays a major role in maintaining rDNA stability, but that Exo1 can substitute for this role when Rad27 is absent. PCNA interacts with Exo1 (*44, 45*). Thus, the partial suppression of rDNA instability by overexpression of *POL30* may indicate that PCNA promotes the recruitment of Exo1 to process Okazaki fragments when Rad27 is absent.

In *Schizosaccharomyces pombe*, meiotic recombination is initiated by DNA double-strand breaks formed by the Spo11 ortholog, Rec12. A previous study demonstrates that cells lacking Rec12 are defective in meiotic recombination, and that deletion of *rad2*^+^/FEN1 from the *rec12*Δ mutant restores meiotic recombination. In the *rad2*Δ *rec12*Δ mutant, meiosis-specific DSBs are undetectable by Southern blotting, leading to a model that cells lacking Rad2 accumulates SSBs or gaps during premeiotic DNA replication, initiating meiotic recombination. Similarly, we observed SSBs in the absence of Rad27, but *rad27*Δ-specific DSBs were undetectable in the rDNA. DSBs may form at the level below the detection limit of Southern blotting, or at random locations, making it difficult to show distinct DSB bands. However, we favor a model that 5’ flap structures are accumulated and converted into 3’ flap structures, to which Rad52 binds and carry out strand invasion into a misaligned rDNA copy, resulting in rDNA copy number changes, independently of DSB formation.

## Data availability

All data are available upon request.

## Acknowledgments

We are grateful to Michael Chan and Eric Alani for the yeast strain and plasmids. We thank Yasuto Murayama at National Institute of Genetics for sharing ideas. We are grateful to Masato Kanemaki and Takuji Iwasato at National Institute of Genetics for the use of Accuri C6 Flow Cytometer and CHEF DR-II System, respectively. We thank the members of the Sasaki laboratory, especially Akemi Mizuguchi for technical assistance.

## Author contributions

M.S. conceived the research, designed the experiments, analyzed the data, and wrote the manuscript. T.Y., N.A., Y.K., and M.S. performed the experiments

## Supplementary Materials

Materials and Methods

### Yeast Strains and Culture Methods

Yeast strains used in this study are derivatives of W303 (*MATa ade2-1 ura3-1 his3-11, 15 trp1-1 leu2-3,112 can1-100*) and are listed in Table S1. Haploid strains in which genes of interest were deleted were constructed by standard one-step gene replacement methods, followed by confirmation of genotypes by PCR. The haploid WT and *cdc9-1* mutants analyzed in Fig. 4A were isolated by tetrad dissection from the ZYY236, a gift from Michael Chen.

Strains carrying plasmids were constructed by standard transformation methods. Yeast cells were grown in Yeast extract-Peptone-Dextrose (YPD) medium (10 g/L yeast extract, 20 g/L peptone, and 20 g/L glucose). For Fig. 5A, cells carrying plasmids were selected in Synthetic complete (SC)-Ura medium [6.7 g/L yeast nitrogen base without amino acids (BD, 291940), 1.92 g/L yeast synthetic drop-out medium supplements without uracil (Sigma, Y1501), 20 g/L glucose]. Agar plates were prepared similarly, with agar added at 20 g/L.

For pulsed-field gel electrophoresis (PFGE) and ERC analyses, 5 mL of appropriate liquid media was inoculated with cells and grown overnight to saturation at 30°C. Cultures were collected (5 × 10^7^ cells/plug), washed twice with 50 mM EDTA (pH 7.5), and cell pellets were stored at - 20°C. For RNA preparation, overnight cultures were diluted in 15 mL of YPAD medium (YPD containing 40 μg/mL adenine sulfate) to OD_600_ = 0.2 and grown until OD_600_ = 0.8. Cultures equivalent to ∼1 × 10^8^ cells were collected, washed twice with ice-cold water, snap-frozen in liquid nitrogen, and stored at -80°C.

For DSB and SSB analyses, *GAL-FOB1* strains were grown overnight in YPD medium at 30°C. Cells were inoculated into ∼200 mL of YP medium supplemented with 40 μg/mL adenine sulfate, 2% raffinose, and one drop of antifoam 204 (Sigma) at a density to reach ∼1 × 10^7^ cells/mL the following morning and grown overnight at 23°C. To arrest cells in G1 phase, α-factor [synthesized by BEX Co. Ltd. (Tokyo, Japan)] was added to cultures at a final concentration of 20 nM. Cells were incubated at 23°C for 3–4 hrs until >80% of the cells were arrested in G1. Then, 20% galactose was added at a final concentration of 2% to induce *FOB1* expression and the cultures were further incubated for 30 min at 23°C. Cells were collected by centrifugation for 5 min at 4°C at 4,500 × *g* and washed twice with YP medium containing 40 μg/mL adenine sulfate and 2% galactose. Cell pellets were resuspended at ∼1 × 10^7^ cells/mL in fresh YP medium containing 40 μg/mL adenine sulfate, 2% galactose, 0.075 mg/mL pronase E (Sigma), and one drop of antifoam 204 (Sigma), then incubated at 23°C. At various time points, 10 mL culture aliquots equivalent to 1 × 10^8^ cells were collected, immediately treated with 1/100 volume of 10% sodium azide, collected, washed twice with 50 mM EDTA pH 7.5, and stored at -20°C.

### Plasmid construction

YCp33 carrying *RAD27* (pMS124), *DNA2* (pMS125), *EXO1* (pMS126), *PIF1* (pMS127), *RNH201* (pMS128), *POL30* (pMS129), and *CDC9* (pMS130) was constructed by Gibson assembly as follows. The genomic regions containing the open reading frame for the gene of interest, along with ∼a few hundred bp upstream and downstream regions were amplified by PCR from the *S. cerevisiae* WT strain of the W303 background, using the following primers: *RAD27*, 5’- TGATTACGCCATTAACAGTGACTTTCGGTG and 5’- CATGCAAGCTTGTCGAAGGCATTACGATGG; *DNA2*-1, 5’- GATTACGCCATCGTAGTTGCTACCTACCAC and 5’- CCTAAAGTCATGACAAGGCTTGGTTCTCCG; DNA2-2, 5’- CTTGTCATGACTTTAGGCAATATCGTACAC and 5’- CATGCAAGCTTGTGTTTTGTGGCTGAGGG; *EXO1*, 5’- ATTACGCCAAAGCTATGTTGGGTGGAACTG and 5’- CTCGCAAAACATATAAAGTTGTCCTC; *PIF1*, 5’- ATTACGCCAAATCAGGACGCTTCAAATGCC and 5’- TGCAAGCTTTTCTATCGAAGGAGGTTCACC; *RNH201*, 5’- ACGCCAAATTGACAGGAAACAAAACTGAG and 5’- AAGCTAACGTAACAGCCTCTCTTCTAG; *POL30*, 5’- ACGCCAAACTTCAGAAAGAAGTTGAGTAGG and 5’- TGCAAGCTATGTGGAAACCCTGAATACCAC; *CDC9*, 5’- TTACGCCATTCCGTAAGACTTGCAATAGG and 5’- AGCTTCTGGATATCTCTCAGTCATATTCTC. YCp33 was linearized by PCR, using the following primers: *RAD27*, 5’- TCACTGTTAATGGCGTAATCATGGTCATAG and 5’- GCCTTCGACAAGCTTGCATGCCTGCAG; *DNA2*-1, 5’- AACTACGATGGCGTAATCATGGTCATAG and 5’- ACAAAACACAAGCTTGCATGCCTGCAG; *EXO1*, 5’- ACATAGCTTTGGCGTAATCATGGTCATAGC and 5’- TATATGTTTTGCGAGAGCTTGCATGCCTGC; *PIF1*, 5’- CGTCCTGATTTGGCGTAATCATGGTCATAG and 5’- TTCGATAGAAAAGCTTGCATGCCTGCAG; *RNH201*, 5’- CCTGTCAATTTGGCGTAATCATGGTCATAG and 5’- AGGCTGTTACGTTAGCTTGCATGCCTGCAG; *POL30*, 5’- TTCTGAAGTTTGGCGTAATCATGGTCATAG and 5’- AGGGTTTCCACATAGCTTGCATGCCTGCAG; *CDC9*, 5’- TCTTACGGAATGGCGTAATCATGGTCATAG and 5’- TGAGAGATATCCAGAAGCTTGCATGCCTGC. These PCR fragments had short homologies at each end and were fused using Gibson Assembly (New England Biolabs) according to the manufacturer’s instructions. Plasmid DNA was sequenced to confirm that it had the proper insertion.

### Fluorescence-activated cell sorter analysis

Approximately 1 × 10^7^ cells were collected at various time points during time course experiments. Cells were fixed with 200 μL of 70% ethanol at -20°C overnight. Cells were collected, resuspended in 200 μL of 50 mM sodium citrate at pH 7.5 containing 0.25 mg/ml RNaseA (Macherey-Nagel), and incubated for 1 hr at 50°C. Then, 100 μL of 50 mM sodium citrate pH 7.5 containing 1 mg/ml Proteinase K (Nacalai) was added and cells were incubated for 1 hr at 50°C. 300 μL of 50 mM sodium citrate pH 7.5 containing 4 μg/ml propidium iodide (Sigma) was added. Cells were sonicated and diluted with 50 mM sodium citrate pH 7.5 containing 4 μg/ml propidium iodide when necessary and were analyzed using a BD Accuri C6 Flow Cytometer (BD Bioscience).

### Genomic DNA preparation

For PFGE, ERC, DSB, and SSB analyses, genomic DNA was prepared in low melting temperature agarose plugs as described previously (*46, 47*). Briefly, collected cells were resuspended in 50 mM EDTA pH 7.5 (33 μL per 5 × 10^7^ cells) and incubated at 42°C. For each plug, 33 μL cell suspension was mixed with 66 μL of solution 1 [0.83% low-melting-point agarose SeaPlaque GTG (Lonza), 170 mM sorbitol, 17 mM sodium citrate, 10 mM EDTA (pH 7.5), 0.85% β-mercaptoethanol, and 0.17 mg/mL Zymolyase 100 T (Nacalai)], poured into a plug mold (Bio-Rad), and placed at 4°C until the agarose was solidified. Plugs were transferred to a 2 mL tube containing solution 2 [450 mM EDTA pH 7.5, 10 mM Tris-HCl pH 7.5, 7.5% β-mercaptoethanol and 10 μg/mL RNaseA (Macherey-Nagel)], and incubated for 1–1.25 hrs at 37°C. Plugs were then incubated overnight at 50°C in solution 3 [250 mM EDTA pH 7.5, 10 mM Tris-HCl pH 7.5, 1% sodium dodecyl sulfate (SDS), and 1 mg/mL proteinase K (Nacalai)]. Plugs were washed four times with 50 mM EDTA (pH 7.5) and stored at 4°C in 50 mM EDTA (pH 7.5).

### PFGE

PFGE was performed as described previously (*46, 47*). Briefly, one-third of a plug was placed on a tooth of the comb, alongside a piece of *Hansenula wingei* chromosomal DNA markers (Bio-Rad). The comb was placed in a gel tray, and 1.0% agarose solution (Pulsed Field Certified Agarose, Bio-Rad) in 0.5× Tris-Borate-EDTA (TBE) (44.5 mM Tris base, 44.5 mM boric acid and 1 mM EDTA pH 8.0) was poured. PFGE was performed on a Bio-Rad CHEF Mapper XA Chiller System or CHEF DR-II System in 2.2 L of 0.5× TBE under the following conditions: 3.0 V/cm for 68 hrs at 14°C, 120° included angle, with initial and final switch times of 300 s and 900 s, respectively. After electrophoresis, DNA was stained with 0.5 μg/mL Ethidium Bromide (EtBr) for 30 min, washed with dH_2_O for 30 min, and then photographed.

### Southern blotting

### Agarose gel electrophoresis

#### ERC assay

The ERC assay was performed as described previously (*47, 48*). Half of the agarose plug was placed on a tooth of the comb. After setting the comb in the gel tray (15 × 25 cm), 300 mL of 0.4% agarose (SeaKem LE Agarose, Lonza) in 1× Tris-acetate-EDTA (TAE) (40 mM Tris base, 20 mM acetic acid, and 1 mM EDTA pH 8.0) was poured into the tray and allowed to solidify. Lambda HindIII DNA marker (600 ng) was loaded in an empty lane. Electrophoresis was performed on a Sub-cell GT electrophoresis system (Bio-Rad) in 1.45 L of 1× TAE at 1.0 V/cm for ∼20–48 hrs at 4°C with buffer circulation. The buffer was changed every ∼24 hrs. DNA was stained with 0.5 μg/mL EtBr for 30 min and then photographed.

#### DSB assay

The DSB assay was performed as described previously (*46, 47*). One-third of an agarose plug was cut and placed in a 2 mL tube. The plug was equilibrated four times in 1 mL of 1× TE by rotating the tube for 15 min at room temperature. The plug was then equilibrated twice in 1 mL of 1× NEBuffer 3.1 (New England Biolabs) by rotating the tube for 30 min at room temperature. After discarding the buffer, the plug was incubated in 160 μL of 1× NEBuffer 3.1 buffer containing 160 units of Bgl II (New England Biolabs) overnight at 37°C. The plug was placed on a tooth of the comb which was set into the gel tray (15 × 25 cm), into which 0.7% agarose solution in 1× TBE was poured; after the gel was solidified, 600 ng of lambda BstEⅡ DNA markers were loaded in an empty lane. Electrophoresis was performed on a Sub-cell GT electrophoresis system (Bio-Rad) in 1.45 L of 1× TBE at 2.0 V/cm for 22 hrs at room temperature with buffer circulation. After electrophoresis, DNA was stained with 0.5 μg/mL EtBr for 30 min and then photographed.

#### SSB assay

One-third of an agarose plug was cut and placed in a 2 mL tube and first digested with Bgl II, as described above. The plug was briefly washed with autoclaved dH_2_O. The plug was then equilibrated twice in 1 mL of 5× alkaline buffer (0.25 M NaOH, 5 mM EDTA [pH 8.0]) by rotating the tube for 15 min at room temperature. The plug was then equilibrated twice in 1 mL of 1× alkaline buffer (0.05 M NaOH, 1 mM EDTA [pH 8.0]) by rotating the tube for 15 min at room temperature. The plug was placed on a tooth of the comb. The agarose solution was prepared by dissolving 3.0 g of SeaKem LE agarose (Lonza) in 270 mL of Milli-Q water by microwave, followed by cooling to 50°C. Then, 30 mL of 10× alkaline buffer was added to the agarose solution and mixed. The comb where the plug was placed was set into the gel tray (15 × 25 cm), and the agarose solution was poured into a gel tray in a cold room, and left until the gel solidified. The comb was removed. DNA markers were prepared in 10 μL containing 1 μg of 1 kb DNA ladder (New England Biolabs), 1 μg of 100 bp DNA ladder (New England Biolabs), 1 μL of 10× alkaline buffer, 1 μL of 0.1% bromophenol blue, and 1 μL of 50% glycerol, which were heat-denatured for 5 min at 70°C, followed by rapid chilling on ice. DNA markers were applied into the well. The gel was run in 1.45 L of 1× alkaline buffer for ∼15 hrs at 4°C at 50 V.

### DNA transfer

After agarose gel electrophoresis, DNA was vacuum-transferred to Nytran SPC (Cytiva) using VacuGene XL (GE Healthcare). Vacuum was applied at 55 cm Hg for 30 min with 0.25 N HCl, for 20 min with denaturation solution (0.5 N NaOH, 1.5 M NaCl), and for 1 hr 40 min–2 hrs with transfer buffer (0.25 N NaOH, 1.5 M NaCl). For SSB assays, transfer was performed at 55 cm Hg for 2 hrs with transfer buffer (0.25 N NaOH, 1.5 M NaCl). After transfer, DNA was fixed to the membrane by soaking the membrane in 300 mL of freshly prepared 0.4 N NaOH for 10 min with gentle shaking, followed by rinsing the membrane with 2× SSC for 10 min. The membrane was subsequently dried and stored at 4°C.

### Probe preparation

Probes were prepared as described previously (*46, 47*). Double-stranded DNA fragments were amplified by PCR. Probe 1 used for ERC and Okazaki fragment analyses was amplified with the primers 5’-CATTTCCTATAGTTAACAGGACATGCC and 5’-AATTCGCACTATCCAGCTGCACTC, and probe 4 used for DSB analysis was amplified with the primers 5’-ACGAACGACAAGCCTACTCG and 5’-AAAAGGTGCGGAAATGGCTG. A portion of the PCR products was gel-purified and used as templates for a second round of PCR with the same primers. The PCR products were then gel-purified. Fifty ng of purified PCR products was used for random priming reactions in the presence of the radiolabeled nucleotide, [α-^32^P]-dCTP (3,000 Ci/mmol, 10 mCi/mL, Perkin Elmer), using Random Primer DNA Labeling Kit (TaKaRa), according to the manufacturer’s instructions. Unincorporated nucleotides were removed using ProbeQuant G-50 Micro Columns (GE Healthcare). Radiolabeled probes were heat-denatured for 5 min at 100°C immediately prior to hybridization to the membrane.

### Hybridization

Southern hybridization was performed as described previously (*46, 47*). The membrane was prewetted with 0.5 M phosphate buffer pH 7.2 and prehybridized for 1 hr at 65°C with 25 mL of hybridization buffer (1% bovine serum albumin, 0.5 M phosphate buffer, pH 7.2, 7% SDS, 1 mM EDTA pH 8.0). After discarding the buffer, the membrane was hybridized overnight at 65°C with 25 mL of hybridization buffer containing heat-denatured probe. The membrane was washed four times for 15 min at 65°C with wash buffer (40 mM phosphate buffer pH 7.2, 1% SDS, 1 mM EDTA pH 8.0) and exposed to a phosphor screen.

### Image analysis

#### ERC assay

Radioactive signals were detected using a Typhoon FLA 9000 (GE Healthcare). The chromosomal rDNA signal was used for normalization of the ERC signals. Thus, the membranes hybridized with radioactive probes were exposed to the phosphor screen for a short time and scanned before signal saturation. Then, membranes were re-exposed to a phosphor screen for several days to achieve optimal signal-to-noise ratios for ERCs. For detection of DIG-labeled probes, chemiluminescent signals were detected after short exposure before signal saturation. Subsequently, membranes were exposed for a longer time to achieve optimal signal-to-noise ratio for ERCs. ERC bands and chromosomal rDNA were quantified from images after long and short exposures, respectively, using Amersham ImageQuant TL ver. 10.2 (Cytiva). ERC levels were calculated as the ratio of ERC signal intensities to chromosomal rDNA signals. When the levels of ERCs were compared between samples loaded on different gels, control DNA plugs were included on each gel for normalization.

#### DSB assay

Membranes were exposed to a phosphor screen for 1–4 days until signals of linear molecules, but not arrested forks, DSBs, or resected DSB signals, reached saturation. The frequency of DSBs and resected DSBs was determined as described previously (*46, 47*). Briefly, in each lane, signal intensities of arrested forks, DSBs, resected DSBs, and the terminal fragment containing the telomere-proximal rDNA repeat and its adjacent non-rDNA fragment were quantified using Amersham ImageQuant TL ver. 10.2 (Cytiva). All signals were normalized to the terminal fragment, and the value at t = 0 min was then subtracted from those at other time points. The DSB frequency was calculated as the ratio of maximum DSB level to maximum arrested fork level in the same time course. The frequency of DSB end resection was determined by dividing the maximum level of resected DSBs by the maximum level of DSBs in the same time course.

### Yeast RNA preparation

RNA was prepared as described previously (*49*), with slight modifications. Collected cells were resuspended in 400 μL of TES (10 mM Tris-HCl pH 7.5, 10 mM EDTA pH 7.5, 0.5% SDS) and 400 μL of acidic phenol by vortexing for approximately 10 s. Cells were incubated at 65°C for 1 hr with occasional vortexing every 15 min. Cell suspensions were then incubated for 5 min on ice and centrifuged at 20,000 × *g* for 10 min at 4°C. The aqueous phase was transferred to a new tube and mixed with an equal volume of acidic phenol by vortexing for 10 s. After 5 min incubation on ice and centrifugation at 20,000 × *g* for 10 min at 4°C, the aqueous phase was transferred to a new tube. RNA was precipitated overnight at -20°C by adding 1/10 volume of 3 M sodium acetate (pH 5.3) and 2.5 volumes of 100% ethanol. The precipitate was collected by centrifugation at 20,000 × *g* for 10 min at 4°C and washed with 70% ethanol. RNA pellets were resuspended in 30 μL of dH_2_O treated with 0.1% diethylpyrocarbonate (DEPC). RNA concentration was determined using Multiskan SkyHigh Microplate Spectrophotometer (Thermo Fisher Scientific). Samples were stored at -80°C.

### Northern blotting

Total RNA (30 μg) was brought up to 7 μL with DEPC-treated dH_2_O and mixed with 17 μL of RNA sample buffer (396 μL of deionized formamide, 120 μL of 10× MOPS buffer [0.2 M MOPS, 50 mM sodium acetate pH 5.2, 10 mM EDTA pH 7.5 in DEPC-treated dH_2_O], and 162 μL of 37% formaldehyde). A total of 1.8 μg of DynaMarker RNA High Markers (BioDynamics Laboratory) were brought up to 4.8 μL with DEPC-treated dH_2_O and mixed with 11.2 μL of RNA sample buffer. Samples were heated at 65°C for 20 min and rapidly chilled on ice. To the samples, 6 μL of 6× Gel Loading Dye (B7025S, New England Biolabs) and 1.5 μL of 2.5 mg/mL EtBr were added. To the size markers, 4 μL of 6× Gel Loading Dye (B7025S, New England Biolabs) and 1 μL of 2.5 mg/mL EtBr were added.

Two 1% agarose gels were prepared by dissolving 1 g of agarose in 73 mL of DEPC-treated dH_2_O. After cooling to 60°C, 17 mL of 37% formaldehyde and 10 mL of 10× MOPS buffer were added. The solution was poured into a gel tray (13 × 12 cm) and allowed to set. Agarose gel electrophoresis was performed on Mini-Sub Cell GT Cell (Bio-Rad) in 650 mL of 1× MOPS buffer at 20 V for 20 min and then at 100 V until the bromophenol blue dye migrated approximately 2/3 of the gel. Gels were photographed and RNA was transferred to Hybond-N+ (GE Healthcare) by standard capillary transfer.

Strand-specific probes were prepared from double-stranded DNA fragments, amplified by PCR and gel-purified. The PCR primers used for probe 2 (against IGS1-R, IGS2-R) were 5’-TCGCCAACCATTCCATATCT and 5’-CGATGAGGATGATAGTGTGTAAGA; for probe 3 (against IGS1-F) were 5’-AGGGAAATGGAGGGAAGAGA and 5’-TCTTGGCTTCCTATGCTAAATCC, and for detecting *TSA1*, 5′-CAAGTTCAAAAGCAAGCTCC and 5′-ACCAATCTCAAGGCTTCGTC. Strand-specific probes were then prepared by linear PCR in a final volume of 20 μL containing 0.2 mM dATP, 0.2 mM dTTP, 0.2 mM dGTP, 50 μL [α-^32^P]-dCTP (3,000 Ci/mmol, 10 mCi/mL, Perkin Elmer), 1.25 units ExTaq (TaKaRa), 1× ExTaq buffer, 50 ng of PCR product as a template, and 10 μM primer (5’-AGTTCCAGAGAGGCAGCGTA for probe 2, 5’-CATTATGCTCATTGGGTTGC for probe 3). PCR was initiated by a denaturation step at 94°C for 3 min, followed by 35 cycles of amplification (96°C for 20 s, 51°C for 20 s, and 72°C for 30 s), and a final step at 72°C for 3 min. Unincorporated nucleotides were removed using ProbeQuant G-50 Micro Columns (GE Healthcare). Radiolabeled probes were heat denatured by incubating for 5 min at 100°C immediately prior to hybridization to the membrane.

Membranes were incubated with 10 mL of ULTRAhyb Ultrasensitive Hybridization Buffer (Thermo Fisher) at 42°C for 1 h. The heat-denatured probe was incubated with the membrane overnight at 42°C. The membrane was rinsed twice with 2× SSC, washed for 15 min at 42°C twice with wash buffer 1 (2× SSC, 0.1% SDS), and washed for 15 min at 42°C twice with wash buffer 2 (0.1× SSC, 0.1% SDS). The membrane was exposed to phosphor screens for several days, and radioactive signals were detected using a Typhoon FLA 7000 (GE Healthcare). Probes were stripped by incubating the membrane with boiled 0.1% SDS by shaking for ∼30 min, rinsed with 2× SSC, and rehybridized with the *TSA1* probe that was prepared as described above. Signals of IGS-F, IGS1-R, IGS2-R, and *TSA1* were quantified using Amersham ImageQuant TL ver. 10.2 (Cytiva). The levels of IGS transcripts were normalized to the *TSA1* signal.

## Quantification and statistical analysis

Statistical analyses were performed using GraphPad Prism, as described in figure legends.

## REFERENCES

1. R. Beroukhim et al., The landscape of somatic copy-number alteration across human cancers. Nature 463, 899–905 (2010).

2. V. Rendo et al., A compendium of Amplification-Related Gain Of Sensitivity genes in human cancer. Nat Commun 16, 1077 (2025).

3. U. Ben-David, A. Amon, Context is everything: aneuploidy in cancer. Nat Rev Genet 21, 44–62 (2020).

4. P. J. Hastings, J. R. Lupski, S. M. Rosenberg, G. Ira, Mechanisms of change in gene copy number. Nat Rev Genet 10, 551–564 (2009).

5. M. Sasaki, J. Lange, S. Keeney, Genome destabilization by homologous recombination in the germ line. Nat Rev Mol Cell Biol 11, 182–195 (2010).

6. R. G. W. Verhaak, V. Bafna, P. S. Mischel, Extrachromosomal oncogene amplification in tumour pathogenesis and evolution. Nat Rev Cancer 19, 283–288 (2019).

7. C. M. Carvalho, J. R. Lupski, Mechanisms underlying structural variant formation in genomic disorders. Nat Rev Genet 17, 224–238 (2016).

8. Y. Hori, C. Engel, T. Kobayashi, Regulation of ribosomal RNA gene copy number, transcription and nucleolus organization in eukaryotes. Nat Rev Mol Cell Biol 24, 414–429 (2023).

9. J. O. Nelson, G. J. Watase, N. Warsinger-Pepe, Y. M. Yamashita, Mechanisms of rDNA Copy Number Maintenance. Trends Genet 35, 734–742 (2019).

10. M. Sasaki, T. Kobayashi, Regulatory processes that maintain or alter ribosomal DNA stability during the repair of programmed DNA double-strand breaks. Genes Genet Syst 98, 103–119 (2023).

11. B. J. Brewer, W. L. Fangman, A replication fork barrier at the 3’ end of yeast ribosomal RNA genes. Cell 55, 637–643 (1988).

12. B. J. Brewer, D. Lockshon, W. L. Fangman, The arrest of replication forks in the rDNA of yeast occurs independently of transcription. Cell 71, 267–276 (1992).

13. T. Kobayashi, M. Hidaka, M. Nishizawa, T. Horiuchi, Identification of a site required for DNA replication fork blocking activity in the rRNA gene cluster in Saccharomyces cerevisiae. Mol Gen Genet 233, 355–362 (1992).

14. M. D. Burkhalter, J. M. Sogo, rDNA enhancer affects replication initiation and mitotic recombination: Fob1 mediates nucleolytic processing independently of replication. Molecular cell 15, 409–421 (2004).

15. T. Kobayashi, T. Horiuchi, P. Tongaonkar, L. Vu, M. Nomura, SIR2 regulates recombination between different rDNA repeats, but not recombination within individual rRNA genes in yeast. Cell 117, 441–453 (2004).

16. T. Weitao, M. Budd, L. L. Hoopes, J. L. Campbell, Dna2 helicase/nuclease causes replicative fork stalling and double-strand breaks in the ribosomal DNA of Saccharomyces cerevisiae. The Journal of biological chemistry 278, 22513–22522 (2003).

17. S. Hosoyamada, M. Sasaki, T. Kobayashi, The CCR4-NOT Complex Maintains Stability and Transcription of rRNA Genes by Repressing Antisense Transcripts. Molecular and cellular biology 40, (2019).

18. J. Houseley, D. Tollervey, Repeat expansion in the budding yeast ribosomal DNA can occur independently of the canonical homologous recombination machinery. Nucleic Acids Res 39, 8778–8791 (2011).

19. S. Ide, K. Saka, T. Kobayashi, Rtt109 prevents hyper-amplification of ribosomal RNA genes through histone modification in budding yeast. PLoS Genet 9, e1003410 (2013).

20. M. Kaeberlein, M. McVey, L. Guarente, The SIR2/3/4 complex and SIR2 alone promote longevity in Saccharomyces cerevisiae by two different mechanisms. Genes Dev 13, 2570–2580 (1999).

21. K. Saka, A. Takahashi, M. Sasaki, T. Kobayashi, More than 10% of yeast genes are related to genome stability and influence cellular senescence via rDNA maintenance. Nucleic Acids Res, (2016).

22. J. L. Stodola, P. M. Burgers, Mechanism of Lagging-Strand DNA Replication in Eukaryotes. Adv Exp Med Biol 1042, 117–133 (2017).

23. L. Zheng, B. Shen, Okazaki fragment maturation: nucleases take centre stage. J Mol Cell Biol 3, 23–30 (2011).

24. L. D. Finger et al., The wonders of flap endonucleases: structure, function, mechanism and regulation. Subcell Biochem 62, 301–326 (2012).

25. L. Zheng et al., Fen1 mutations result in autoimmunity, chronic inflammation and cancers. Nat Med 13, 812–819 (2007).

26. M. Sasaki, T. Kobayashi, Transcription near arrested DNA replication forks triggers ribosomal DNA copy number changes. Nucleic Acids Res 53, (2025).

27. A. S. Ivessa, J. Q. Zhou, V. A. Zakian, The Saccharomyces Pif1p DNA helicase and the highly related Rrm3p have opposite effects on replication fork progression in ribosomal DNA. Cell 100, 479–489 (2000).

28. M. R. Lieber, The FEN-1 family of structure-specific nucleases in eukaryotic DNA replication, recombination and repair. Bioessays 19, 233–240 (1997).

29. Y. Xie et al., Identification of rad27 mutations that confer differential defects in mutation avoidance, repeat tract instability, and flap cleavage. Molecular and cellular biology 21, 4889–4899 (2001).

30. M. C. Negritto, J. Qiu, D. O. Ratay, B. Shen, A. M. Bailis, Novel function of Rad27 (FEN-1) in restricting short-sequence recombination. Molecular and cellular biology 21, 2349–2358 (2001).

31. T. Kobayashi, A. R. Ganley, Recombination regulation by transcription-induced cohesin dissociation in rDNA repeats. Science 309, 1581–1584 (2005).

32. C. C. Liu et al., Flap endonuclease Rad27 cleaves the RNA of R-loop structures to suppress telomere recombination. Nucleic Acids Res 51, 4398–4414 (2023).

33. J. Houseley, K. Kotovic, A. El Hage, D. Tollervey, Trf4 targets ncRNAs from telomeric and rDNA spacer regions and functions in rDNA copy number control. EMBO J 26, 4996–5006 (2007).

34. M. Yokoyama, M. Sasaki, T. Kobayashi, Spt4 promotes cellular senescence by activating non-coding RNA transcription in ribosomal RNA gene clusters. Cell Rep 42, 111944 (2023).

35. R. D. Kolodner, G. T. Marsischky, Eukaryotic DNA mismatch repair. Curr Opin Genet Dev 9, 89–96 (1999).

36. L. H. Johnston, K. A. Nasmyth, Saccharomyces cerevisiae cell cycle mutant cdc9 is defective in DNA ligase. Nature 274, 891–893 (1978).

37. D. J. Smith, I. Whitehouse, Intrinsic coupling of lagging-strand synthesis to chromatin assembly. Nature 483, 434–438 (2012).

38. H. Debrauwere, S. Loeillet, W. Lin, J. Lopes, A. Nicolas, Links between replication and recombination in Saccharomyces cerevisiae: a hypersensitive requirement for homologous recombination in the absence of Rad27 activity. Proceedings of the National Academy of Sciences of the United States of America 98, 8263–8269 (2001).

39. D. X. Tishkoff, N. Filosi, G. M. Gaida, R. D. Kolodner, A novel mutation avoidance mechanism dependent on S. cerevisiae RAD27 is distinct from DNA mismatch repair. Cell 88, 253–263 (1997).

40. R. Ayyagari, X. V. Gomes, D. A. Gordenin, P. M. Burgers, Okazaki fragment maturation in yeast. I. Distribution of functions between FEN1 AND DNA2. The Journal of biological chemistry 278, 1618–1625 (2003).

41. H. I. Kao, J. Veeraraghavan, P. Polaczek, J. L. Campbell, R. A. Bambara, On the roles of Saccharomyces cerevisiae Dna2p and Flap endonuclease 1 in Okazaki fragment processing. The Journal of biological chemistry 279, 15014–15024 (2004).

42. M. Levikova, P. Cejka, The Saccharomyces cerevisiae Dna2 can function as a sole nuclease in the processing of Okazaki fragments in DNA replication. Nucleic Acids Res 43, 7888–7897 (2015).

43. M. Kahli, J. S. Osmundson, R. Yeung, D. J. Smith, Processing of eukaryotic Okazaki fragments by redundant nucleases can be uncoupled from ongoing DNA replication in vivo. Nucleic Acids Res 47, 1814–1822 (2019).

44. X. Chen, S. C. Paudyal, R. I. Chin, Z. You, PCNA promotes processive DNA end resection by Exo1. Nucleic Acids Res 41, 9325–9338 (2013).

45. A. Cheruiyot et al., Poly(ADP-ribose)-binding promotes Exo1 damage recruitment and suppresses its nuclease activities. DNA repair 35, 106–115 (2015).

46. M. Sasaki, T. Kobayashi, Ctf4 Prevents Genome Rearrangements by Suppressing DNA Double-Strand Break Formation and Its End Resection at Arrested Replication Forks. Molecular cell 66, 533–545 e535 (2017).

47. M. Sasaki, T. Kobayashi, Gel Electrophoresis Analysis of rDNA Instability in Saccharomyces cerevisiae. Methods Mol Biol 2153, 403–425 (2021).

48. M. Goto, M. Sasaki, T. Kobayashi, The S-Phase Cyclin Clb5 Promotes rRNA Gene (rDNA) Stability by Maintaining Replication Initiation Efficiency in rDNA. Molecular and cellular biology 41, (2021).

49. T. Iida, T. Kobayashi, RNA Polymerase I Activators Count and Adjust Ribosomal RNA Gene Copy Number. Molecular cell 73, 645–654 e613 (2019).

