## Supplemental Table 1 for "Rad27/FEN1 prevents accumulation of unprocessed Okazaki fragments and ribosomal DNA copy number changes"

**Supplementary Table 1. *S. cerevisiae* strains used in this study**

| Name | Genotype |
|---|---|
| MSY275 | *MATa* |
| MSY360 | *MATa, NatNT2-GALL-FOB1, bar1::LEU2* |
| MSY409 | *MATa, fob1::LEU2* |
| MSY937 | *MATa, NatNT2-GALL-FOB1, bar1::LEU2, sir2Δ::hphMX, hmlΔ::kanMX* |
| MSY1645 | *MATa, sir2Δ::hphMX* |
| MSY1648 | *MATa, rad27Δ::kanMX* |
| MSY1651 | *MATa, sir2Δ::hphMX, rad27Δ::kanMX* |
| MSY1654 | *MATa, fob1::LEU2, sir2Δ::hphMX* |
| MSY1657 | *MATa, fob1::LEU2, rad27Δ::kanMX* |
| MSY1660 | *MATa, fob1::LEU2, sir2Δ::hphMX, rad27Δ::kanMX* |
| MSY1636 | *MATa, din7Δ::kanMX* |
| MSY1638 | *MATa, yen1Δ::kanMX* |
| MSY1640 | *MATa, exo1Δ::kanMX* |
| MSY1665 | *MATa, RAD27::rad27-G240D::LEU2* |
| MSY1674 | *MATa, RAD27::LEU2* |
| MSY1680 | *MATa, RAD27::rad27-324::LEU2* |
| MSY1684 | *MATa, RAD27::rad27-G67S::LEU2* |
| MSY1687 | *MATa, RAD27::rad27-E158D::LEU2* |
| MSY1790 | *MATa, NatNT2-GALL-FOB1, bar1::LEU2, rad27Δ::kanMX* |
| MSY1898 | *MATa, rad2Δ::kanMX* |
| ZYY236 | *MATa/α, est2Δ::URA3/EST2*, *cdc9-1::kanMX/CDC9* |

All strains are derivatives of W303, which is *ade2-1, ura3-1, his3-11, 15, trp1-1, leu2-3, 112, can1-100*, and *RAD5*.
